# Behavioral and biochemical effects of ethanol withdrawal in zebrafish

**DOI:** 10.1101/201368

**Authors:** Suianny Nayara da Silva Chaves, Gabriel Felício Rocha, Bruna Patrícia Dutra Costa, Witallo Etevaldo Araújo de Oliveira, Monica Gomes Lima, Diógenes Henrique de Siqueira Silva, Caio Maximino

**Affiliations:** Laboratório de Neurociências e Comportamento, Instituto de Estudos em Saúde e Biológicas, Universidade Federal do Sul e Sudeste do Pará; Escola Estadual de Ensino Médio Anísio Teixeira; Laboratório de Neurofarmacologia e Biofísica, Campus VIII, Universidade do Estado do Pará; Programa de Pós-Graduação em Biodiversidade e Biotecnologia – Rede BIONORTE; Programa de Pós-Graduação em Neurociências e Comportamento, Universidade Federal do Pará

**Keywords:** Anxiety, *Danio rerio*, Ethanol withdrawal, Catalase activity, Epileptic seizures.

## Abstract

Chronic alcohol use induces adaptations and toxicity that can induce symptoms of anxiety, autonomic hyperarousal, and epileptic seizures when alcohol is removed (withdrawal syndrome). Zebrafish has recently gained wide attention as a behavioral model to study the neurobehavioral effects of acute and chronic alcohol use, including withdrawal. The literature, however, is very contradictory on findings regarding withdrawal effects, with some studies reporting increased anxiety, while others report no effect. A meta-analytic approach was taken to find the sources of this heterogeneity, and ethanol concentration during exposure and exposure duration were found to be the main sources of variation. A conceptual replication was also made using continuous exposure for 16 days in waterborne ethanol (0.5%) and assessing anxiety-like behavior in the light/dark test after 60 min withdrawal. Withdrawal was shown to reduce preference for darkness, consistent with decreased anxiety, but to increase risk assessment, consistent with increased anxiety. Animals were also subjected to the withdrawal protocol and injected with pilocarpine in a sub-convulsive dose to assess susceptibility to epileptic seizure-like behavior. The protocol was sufficient to increase susceptibility to epileptic seizure-like behavior in animals exposed to ethanol. Finally, withdrawal also decreased catalase activity in the brain, but not in the head kidney, suggesting mechanisms associated with the behavioral effects of ethanol withdrawal.

## 1. Introduction

Chronic alcohol (ethanol, EtOH) use produces adaptations and toxicity that can lead to tolerance and dependence, manifested as physical and mental distress when EtOH is removed (withdrawal); symptoms of ethanol withdrawal include anxiety, insomnia, and autonomic hyperarousal (Krystal and Tabakoff, 2002). In more serious conditions, patients presenting this EtOH withdrawal syndrome can present perceptive changes, agitation, mental confusion, significant increases in autonomic arousal, and epileptic seizures (Gatch and Lal, 2001; Krystal and Tabakoff, 2002). The most serious condition involves *delirium tremens* and death by hyperthermia, cardiac arrhythmia, and complications from withdrawal-induced epileptic seizures (Longo and Schuckit, 2014). These symptoms also present with a typical time course, with marked signs of anxiety appearing as early as 6 hours after cessation of alcohol consumption, and epileptic seizures appearing from 12 to 48 hours after withdrawal (Trevisan et al., 1998). Since withdrawal symptoms are usually reduced after EtOH consumption, EtOH dependence can be maintained by negative reinforcement (Koob and Le Moal, 2008), and therefore investigating these motivational mechanisms could open new avenues for the treatment of EtOH consumption-related disorders.

In animal models, EtOH withdrawal changes the excitability of neurons located in brain regions associated with defensive behavior, anxiety, and fear (Bonassoli et al., 2011; Chakravarty and Faingold, 1998; Long et al., 2007; Yang et al., 2003, 2002, 2001). Moreover, EtOH withdrawal also dysregulates the activity of the hypothalamus-pituitary-adrenal axis that modulates behavioral and endocrine responses to stress (Rasmussen et al., 2002). It makes sense, then, that the majority of animal models of EtOH withdrawal are focused on anxiety-like behavior, with consistent effects observed in rodent models such as the elevated plus-maze, light/dark box, social interaction test, and a drug discrimination assay using pentylenetetrazole (Gatch and Lal, 2001).

Zebrafish (*Danio rerio*) is increasingly being considered as useful model organisms for studying both behavioral genetics and behavioral neuroscience, neuropsychopharmacology, and neurotoxicology (Bonan and Norton, 2015; Kalueff et al., 2012; Norton and Bally-Cuif, 2010; Shams et al., 2018; Stewart et al., 2015). The main advantages associated with this species are its low cost of acquisition and upkeep, ease handling, short lifespan, and readiness of reproduction in laboratory environments (Gerlai, 2014; Kalueff et al., 2014). The relatively high degree of genetic, neural, and endocrine homology with rodents and human beings is also cited as an advantage (Kokel and Peterson, 2008). Zebrafish is also a good model for studying anxiety and stress, with well-validated assays for novelty- and conflict-induced anxiety, social interaction, and antipredatory behavior (Gerlai, 2010; Maximino et al., 2010; Oliveira, 2013). Importantly for EtOH withdrawal, epileptic seizure-like behaviors were also characterized in the species (Hortopan et al., 2010), although not yet in a context of EtOH withdrawal.

Zebrafish anxiety-like behavior has been used to demonstrate the effects of drug withdrawal, including cocaine (López-Patiño et al., 2008a, 2008b), morphine (Cachat et al., 2010; Khor et al., 2011; Wong et al., 2010), and EtOH (Tran et al., 2016). In the last case, the literature is inconsistent, with some studies (e.g., Tran et al., 2015) reporting significant effects of withdrawal on anxiety-like behavior, while others (e. g., Cachat et al., 2010) were unable to detect effects of EtOH withdrawal. Procedural differences, such as assay type, strain, concentration during exposure, or withdrawal duration, could be responsible for this difference.

One important consistency that is found in the literature regards results using the light/dark test. For example, Benneh et al. (2016) and Mathur and Guo (2011) found no effect of withdrawal on dark preference, while Holcombe et al. (2013) found that zebrafish subjected to EtOH withdrawal reversed their preference, spending more time in the white compartment instead of the black compartment. The light/dark test is conceptually different from the novel tank test (one of the most commonly used assays in zebrafish withdrawal research) in that an approach-avoidance conflict appears to underline behavior in the light/dark test, while escape from the top appears to motivate behavior in the novel tank test (Maximino et al., 2012); a recent metanalysis (Kysil et al, 2017) also suggested that the light/dark test is more sensitive to pharmacological treatments than the novel tank test. The discrepancies in the literature regarding the effects of EtOH withdrawal on both tests could represent different underlining neurobiological bases, or methodological differences.

In addition to these behavioral endpoints, neurochemical analyses and oxidative stress assays were also reported in zebrafish models of EtOH withdrawal (Müller et al., 2017; Tran et al., 2015b). After exposure to 0.5% EtOH for 22 days, increases in brain levels of dopamine, serotonin, and aspartate were observed with 60 min withdrawal (Pan et al., 2012). After exposure for 8 days to 1% EtOH (for 20 min per day), decreases in superoxide dismutase and catalase activity and consequent increases in oxidative stress were observed (Müller et al., 2017). While a mechanistic explanation is still lacking, these results are congruent with the hyperexcitability and EtOH-induced neurotoxicity observed in other models (Krystal and Tabakoff, 2002; Zenki et al., 2014).

The aim of the present work is to study the heterogeneity of effects of EtOH withdrawal on zebrafish anxiety-like behavior in the literature, by applying meta-analytical techniques. Moreover, the present work attempts a conceptual replication of findings on EtOH withdrawal by assessing the effects of a withdrawal protocol on behavior in the light/dark test. It also attempts to expand the range of endpoints used in withdrawal research by using a chemically-induced epileptic seizure-like behavior model with sub-convulsive doses. Finally, the present work also attempted to replicate previous research on the effects of EtOH withdrawal on the activity of the enzyme catalase in the brain and head kidney. This manuscript is a complete report of all the studies performed to test the effect of ethanol withdrawal on anxiety-like and convulsive-like behavior in zebrafish. We report how we determined our sample size, all data exclusions (if any), all data transformations, and all measures in the study (Simmons et al., 2012).

## 2. Methods

### 2.1 Systematic review and metanalysis

The protocol for the meta-analysis was pre-registered in the CAMARADES-NC3Rs Preclinical Systematic Review & Meta-analysis Facility (SyRF) database (https://drive.google.com/file/d/0B7Z0eAxKc8ApUjcyQjhwVnFjRFE/view?usp=sharing). No modification from the pre-registered protocol was made. Article with the descriptors ‘ethanol withdrawal’ and ‘zebrafish’ were searched for in PubMed (https://www.ncbi.nlm.nih.gov/pubmed), using a search filter optimized to finding studies on animal experimentation on PubMed (Hooijmans et al., 2010). Bibliographic data (including DOI, publication date, title, and abstract) from the studies identified in the systematic review were exported to a spreadsheet. Each article from the list was reviewed in four levels of detail (title, abstract, full text, and a detailed revision of the experimental design) in order to determine its eligibility to meta-analysis. Following Mohammad et al. (2016), studies should include (1) primary behavioral data obtained in tests for anxiety-like behavior in zebrafish (light/dark test, novel tank test, antipredator responses, shoaling responses); (2) reporting of appropriate controls; and (3) reporting of at least sample sizes and summary statistics (central tendency and dispersion measures) for control and withdrawal groups. While the light/dark and novel tank tests, as well as antipredator responses, straightforwardly represent anxiety-like behavior, shoaling has been included because it is sensitive to manipulations which increase anxiety and/or fear in zebrafish (Green et al., 2012), suggesting a defensive component. When an experiment evaluated the effects of drugs or other interventions on withdrawal syndrome-like effects, only control and EtOH withdrawal groups were considered, and data on intervention effects was not analyzed; e.g., while Pittman and Hylton (2015) assessed the effects of fluoxetine and ketamine on withdrawal-induced anxiogenesis, these effects were not assessed in the metanalysis. Possible confounds in relation to the role of development were reduced by excluding studies which were not performed on adult fish.

The following data were extracted from each included study: identification (DOI, authors, publication year); strain/phenotype; concentration of ethanol during exposure; duration of EtOH treatment; duration of withdrawal; behavioral test that was used; means and standard deviations, as well as a test statistics and degrees of freedom; and sample sizes (*N*) for each group. Data, which were represented graphically, were extracted from figures using PlotDigitizer (http://plotdigitizer.sourceforge.net/). When multiple dependent variables were reported, only the primary endpoint was used (time on white, time on bottom, inter-fish distance, and distance from stimulus). While there is considerable variation in the effects of interventions on these tasks – and indeed variables such as erratic swimming or freezing can be more sensitive to certain treatments –, the literature usually refers to time on white and time on bottom as the primary endpoints. Distance from stimulus was used as a primary endpoint for the shoaling and antipredator behavior experiments made by Robert Gerlai’s group, given that it indicates increased shoaling (decreased distance to shoal stimulus) or predator avoidance (increased distance to predator stimulus). All estimates were transformed to standardized mean differences (SMD), corrected for its positive bias (Hedges and Olkin, 1985), with unbiased estimates of sampling variances and confidence intervals at the 95% level, *I*² and τ² heterogeneity values, and p-values using a mixed-effects model, with concentration, exposure duration, and withdrawal duration used as moderators. Differently from the other endpoints, decreased distances to the shoal stimulus or decreased inter-fish distances indicate less anxiety, and the SMDs for these cases were transformed by multiplying by -1 (Vesterinen et al., 2014). Influential case diagnostics was made by inspecting plots for externally standardized residues, DFFITS values, Cook’s distances, covariance ratios, estimates of τ*²* and test statistics for residual heterogeneity when each study is removed in turn, hat values, and weights for each study included in the analysis. Publication bias was assessed by inspection of a contour-enhanced funnel plot, with contours at the 90%, 95% and 99% confidence intervals. Moreover, funnel plot asymmetry was analyzed using a meta-regression test, with total samples size as predictor (Egger et al., 1997). Observed power for each study was calculated based on effect sizes, sample sizes and standard deviations, and fitted against SMDs by a generalized additive model with Gaussian curve family and identity link function. Finally, a sensitivity analysis was performed by adding study quality, assessed using SYRCLE’s Risk of Bias (RoB) tool (Hooijmans et al., 2014), in the meta-regression model. The meta-analysis was made with the metafor package from R (Viechtbauer, 2010).

### 2.2. Effects of EtOH withdrawal on anxiety-like behavior and epileptic seizure-like behavior susceptibility in zebrafish

#### 2.1.1. Animals, housing, and baseline characteristics

Outbred populations were used due to their increased genetic variability, decreasing the effects of random genetic drift which could lead to the development of uniquely heritable traits (Parra et al., 2009; Speedie and Gerlai, 2008). Thus, the animals used in the experiments are expected to better represent the natural populations in the wild. Twenty-four-adult zebrafish from the wildtype strain (longfin phenotype) were used in this experiment. Animals were bought from a commercial seller, and arrived in the laboratory with 3 months of age, approximately (standard length = 13.2 ± 1.4 mm), and were quarantined for two weeks; the experiment begun when animals had an approximate age of 4 months (standard length = 23.0 ± 3.2 mm). Animals were kept in mixed-sex tanks during acclimation, with an approximate ratio of 50 male: 50 female. The breeder was licensed for aquaculture under Ibama’s (Instituto Brasileiro do Meio Ambiente e dos Recursos Naturais Renováveis) Resolution 95/1993. Animals were group-housed in 40 L tanks, with a maximum density of 25 fish per tank, for at least 2 weeks before experiments begun. Tanks were filled with non-chlorinated water at room temperature (28°C) and a pH of 7.0-8.0. Lighting was provided by fluorescent lamps in a cycle of 14-10 hours (LD), according to standards of care for zebrafish (Lawrence, 2007). Water quality parameters were as follow: pH 7.0-8.0; hardness 100-150 mg/L CaCO3; dissolved oxygen 7.5-8.0 mg/L; ammonia and nitrite < 0.001 ppm. All manipulations minimized their potential suffering of animals, and followed Brazilian legislation (Conselho Nacional de Controle de Experimentação Animal - CONCEA, 2017). Animals were used for only one experiment and in a single behavioral test, to reduce interference from apparatus exposure.

#### 2.2.2. Ethanol exposure and withdrawal

The EtOH exposure and withdrawal regimen was adapted from Gerlai et al. (2009). In brief, animals were exposed to increasing EtOH concentrations (0.125%-0.5%), dispersed on the tank water, for 16 days, therefore decreasing mortality that is associated with prolonged exposure to EtOH in zebrafish. Concentrations doubled every four days, reaching a final concentration of 0.5% (v/v). Animals were kept in this final concentration for 4 days. Another group was exposed to the same manipulation, without EtOH exposure; animals were randomly allocated to treatments *via* generation of random numbers using the randomization tool in http://www.randomization.com/. Caretakers were blinded for treatment. After treatment, animals were transferred to a tank with system water for 60 min (withdrawal stage). All animals were included in the experiments, and no gross physical abnormalities were observed during the exposure period. To control for effects of chronic EtOH exposure that are not attributable to withdrawal, two additional groups of 8 animals were exposed as above and transferred, without an withdrawal stage, to the light/dark tank.

#### 2.2.3. Light/dark test

After EtOH withdrawal, animals (10 animals in the control group, 14 in the withdrawal group) were individually tested in a tank (15 cm height X 10 cm width X 45 cm length) that was divided in half into a black compartment and a white compartment. The tank was made of matte acrylic. The apparatus was illuminated from above by two fluorescent 25 W lamps, which produced an average of 270 lumens above the tank (light levels were measured using a hand photometer). The tank contained sliding doors that defined a central compartment in which the animal was positioned for a 3-min acclimation. After this stage, the sliding doors were removed, allowing the animal to freely explore the apparatus for 15 min.

The order with which animals were exposed to the tank was randomized and balanced across treatments. Experimenters were blinded to treatment. Videos were manually transcribed by two observers, blinded to treatment, using X-Plo-Rat (https://github.com/lanec-unifesspa/x-plo-rat). The following variables were analyzed: time on the white compartment (s); transitions to white; total locomotion on white (number of virtual 4.5 cm² squares crossed by the animal in the compartment); mean duration of entries in the white compartment (total duration divided by the number of transitions); time freezing (s); number of erratic swimming events; time in thigmotaxis (s); number of risk assessment events. Operational definitions of these endpoints can be found in Table 1.

Data were analyzed via approximative two-sample Fisher-Pitman permutation tests with 10.000 Monte-Carlo re-samplings. The data analyst was blinded to treatment by cell scrambling; after analysis, data was unblinded. Data are presented using individual dot plots combined with summaries of mean and bootstrapped confidence intervals at 95% level. Standardized mean differences with unbiased variance estimates were calculated using the R package ‘metafor’. *Post-hoc* (observed) power was calculated based on an approximation of the T-test, with two-tailed hypotheses. All analyses and graphs were made using R version 3.3.0 and packages ‘ggplot2’ and ‘coin’.

**Table 1.** Operational definitions for behavioral endpoints assessed in the light/dark test. Whenever available, codes used in the Zebrafish Behavioral Catalog (Kalueff et al., 2013) are provided.

| Behavioral endpoint | Definition |
| --- | --- |
| Freezing | Time spent immobile, with the exception of opercular and eye movements, in seconds, observed in the white compartment (ZBC 1.68) |
| Erratic swimming | Sharp changes in direction or velocity of swimming, associated with repeated fast acceleration bouts, observed in the white compartment (ZBC 1.51) |
| Thigmotaxis | Percentage of time swimming in a distance of 2 cm or less from the white compartment's walls (ZBC 1.173) |
| Risk assessment | Fast (< 1 second) entries in the white compartment, followed by re-entry in the black compartment, or partial entries in the white compartment (i.e., the pectoral fin does not cross the midline) |

#### 2.2.4. Pilocarpine-induced epileptic seizure-like behaviors

Immediately after the light/dark test, animals were injected with a dose of 150 mg/kg of pilocarpine. This dose has been shown to be insufficient to produce clonic and tonic-clonic epileptic seizure-like behavior in adult animals (Pinto, 2015). Fifteen minutes after injection, animals were individually transferred to 1.5 L tanks, and filmed for 15 min to analyze the profile of epileptic seizure-like behavior. Epileptic seizure-like behaviors were scored according to Mussulini et al. (2013), as in Table 2.

**Table 2.**
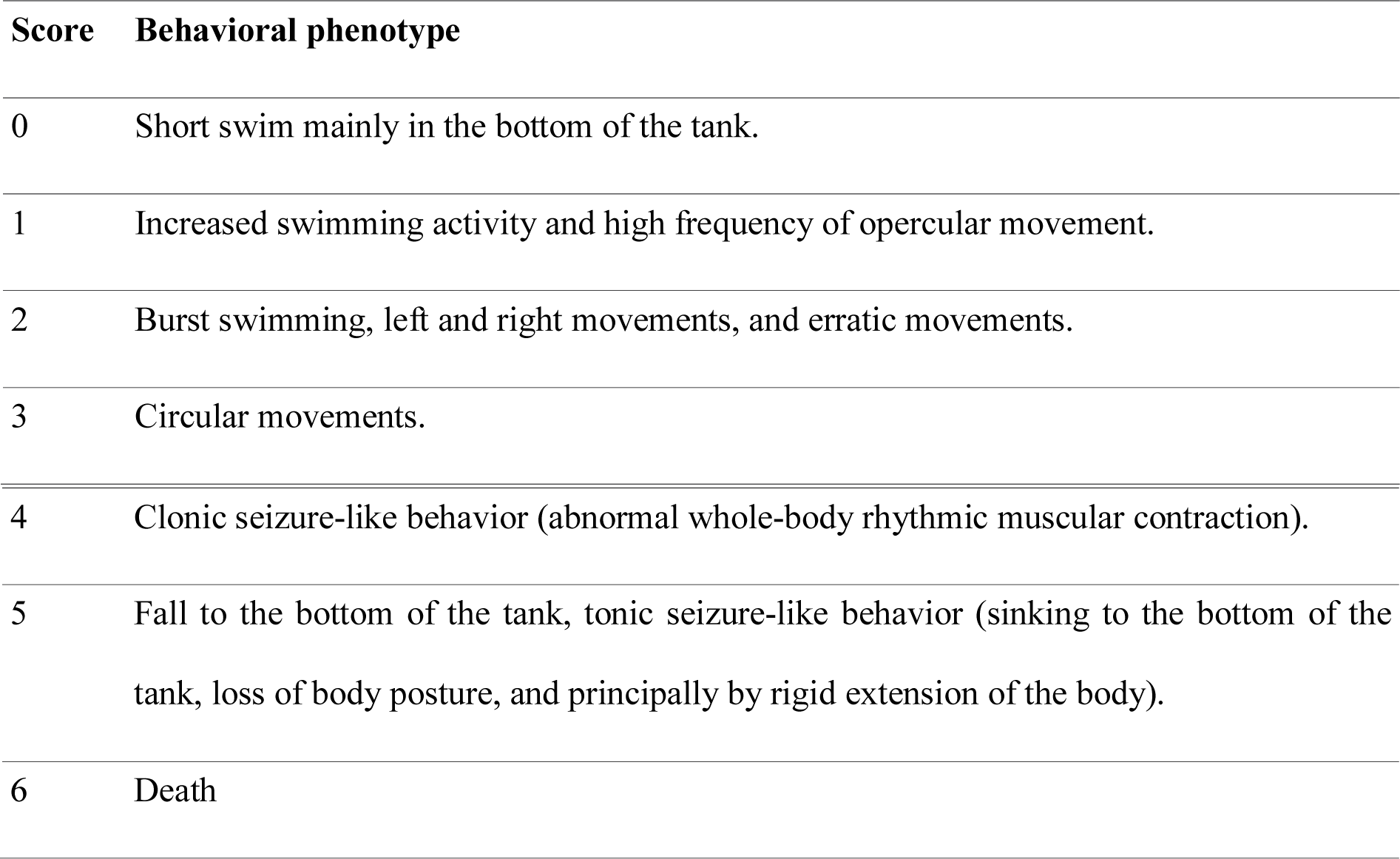
Behavioral phenotypes scored in the pilocarpine-induced seizure model. Scores were based on Mussulini et al. (2013).

| Score | Behavioral phenotype |
| --- | --- |
| 0 | Short swim mainly in the bottom of the tank. |
| 1 | Increased swimming activity and high frequency of opercular movement. |
| 2 | Burst swimming, left and right movements, and erratic movements. |
| 3 | Circular movements. |
| 4 | Clonic seizure-like behavior (abnormal whole-body rhythmic muscular contraction). |
| 5 | Fall to the bottom of the tank, tonic seizure-like behavior (sinking to the bottom of the tank, loss of body posture, and principally by rigid extension of the body). |
| 6 | Death |

Score 4 is the minimum behavioral phenotype that can be considered epileptiform (Mussulini et al., 2013); therefore, the latency to reach Score 4 was considered as the main endpoint for epileptic seizure-like behavior. Latencies were analyzed by fitting a Kaplan-Meier model to survival curves, using the R package ‘survival’.

### 2.3. Effects of EtOH withdrawal on catalase activity

A separate group of 12 animals were used in this experiment. Animals were subjected to the exposure and withdrawal regimen described in 2.2.1. Sixty minutes after withdrawal, animals were euthanized in cold water followed by spinal section, and their brains and head kidneys were dissected. Catalase activity in those organs was measured using the rate of disappearance of H_2_O_2_ spectrophotometrically, following the method described by Aebi (1984). Within-laboratory validation yielded a linearity of r² = 0.9849, and intermediary repeatability of 0.2364 (IC95% [0.0042, 0.4686]; Horwitz ratio). Enzyme activity was corrected by protein levels, quantified by the Bradford method. Differences between groups were analyzed using Approximative Two-Sample Fisher-Pitman Permutation Tests.

## 3. Results

### 3.1. Metanalysis

To evaluate the effect of EtOH withdrawal on zebrafish anxiety-like behavior, we applied a mixed-effects meta-regression model on the results from the systematic review. Characteristics from the articles found in the systematic review can be found in Table 3; raw data and analysis scripts for this metanalysis can be found in our GitHub repository (https://github.com/lanec-unifesspa/etoh-withdrawal/tree/master/metanalysis). EtOH concentrations ranged from 0.25% to 3% v/v (median 1%); exposure durations ranged from 7 to 63 days (median 14 days), and withdrawal durations ranged from 1 to 1.512 h (median 48 h). In general, a high risk of bias was observed, since most studies did not report blinding or random allocation (https://github.com/lanec-unifesspa/etoh-withdrawal/blob/master/metanalysis/etoh-withdrawal-metanalysis-rob.csv).

Behavioral test, strain/phenotype, EtOH concentration, exposure duration, and withdrawal duration were used as moderators. Results from this analysis are presented in the forest plot found in Figure 1A. Residual heterogeneity was estimated as τ*²* = 0.1667, significantly high (QE[_df= 11_] = 22.6338, *p* = 0.0199), suggesting that although about 88% of the total heterogeneity can be explained by including the five moderators in the model, other factors might influence the effect. After applying a permutation test with 1.000 replications, a significant effect of exposure duration (*p* = 0.029) and EtOH concentration (*p* = 0.001) were found, but the other mediators did not affect withdrawal-like behavior (Figures 1B**-1D**). The contour-enhanced funnel plot (Figure 1E) suggest that most studies failed to reach statistical significance, with only one study falling in the 95% CI range, and one in the 99% CI range. Egger’s test on this funnel plot did not suggest publication bias (t[_df= 10_] = 0.439, *p* = 0.67). An analysis of influential observations suggests that the Mathur and Guo (2011) study on the effects on the NTT with the longer withdrawal duration and the Dewari et al. (2016) study produced most residual heterogeneity (**Figure S1**). Absolute SMDs were significantly explained by observed power (slope = 1.2819, *p* = 0.0174; **Figure S2**). Finally, sensitivity analysis suggested that study quality did not influence results, as including RoB scores in the model did not improve fit (model without RoB: AIC = 46.4816, BIC = 50.88=585; model with RoB: AIC = 47.2908, BIC = 50.9218).

**Figure 1.**
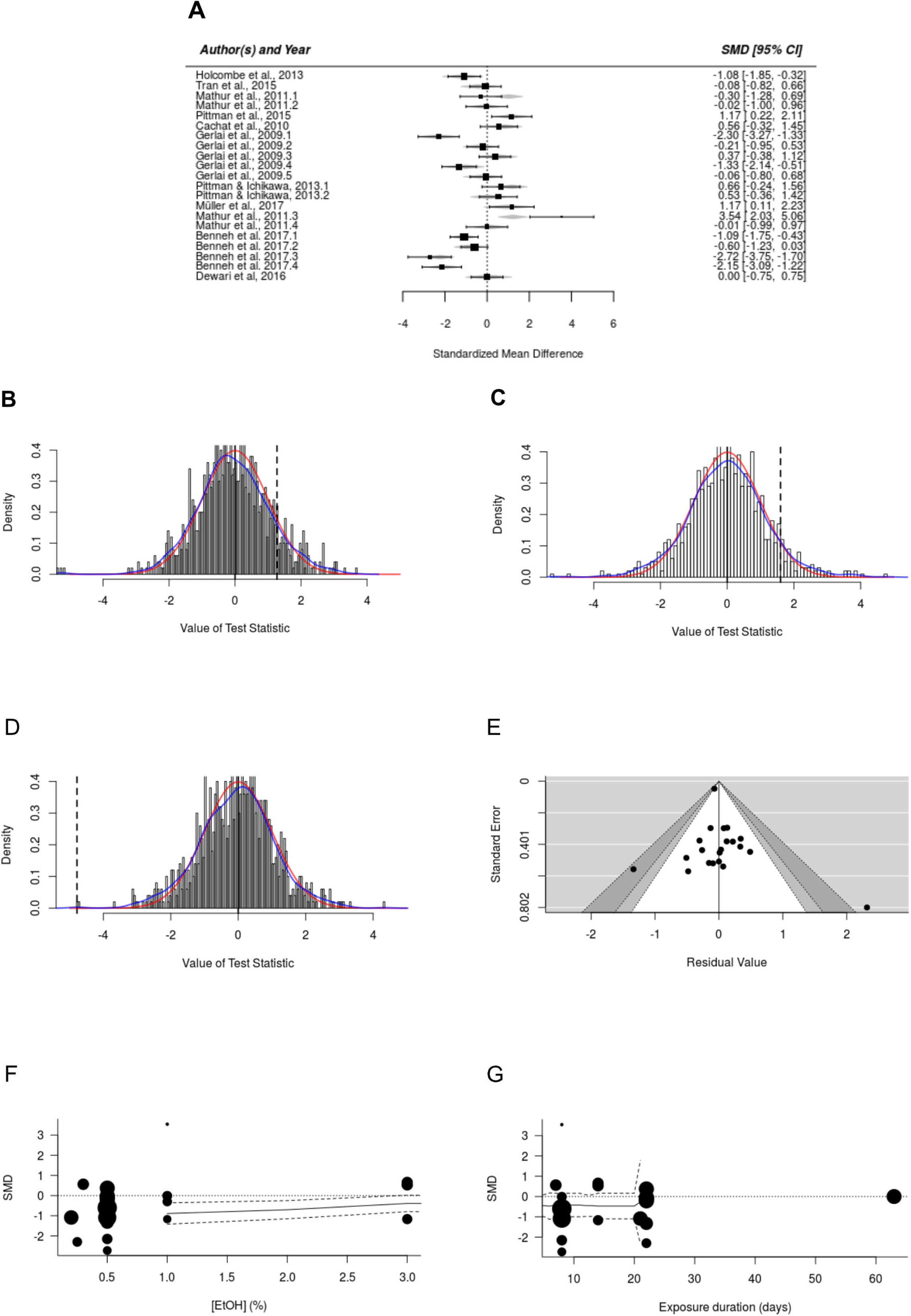
Metanalysis of EtOH withdrawal experiments in zebrafish reveal a significant increase in anxiety-like behavior, but high heterogeneity driven by methodological differences. (A) Forest plot showing the results of 16 studies examining the effect of ethanol withdrawal on zebrafish anxiety-like behavior. The figure shows the standardized mean difference (SMD) between control and withdrawal-exposed groups with corresponding 95% confidence intervals in the individual studies, based on a mixed-effects model. A negative standardized mean difference (SMD) corresponds to decreased anxiety-like behavior, while a positive SMD corresponds to increased anxiety-like behavior after ethanol withdrawal. Studies are ordered by total degrees of freedom. (B-D) Permutation distribution of the test statistic for the mediators: behavioral test (B), ethanol concentration during exposure (C), exposure duration, in days (D), withdrawal duration, in hours (E), and strain/phenotype (F). Distributions were based on a permutation test with 1,000 replications. The blue contour represents kernel density estimates of the permutation distributions; the red curve represents the standard normal density; the full line represents the null hypothesis of no difference; and the dashed line represents the observed values of test statistics. (G) Contour-enhanced funnel plot of meta-analysis. Estimated standardized mean differences were plotted against precision (1/standard error) were A negative estimate corresponds to decreased anxiety-like behavior, while a positive estimate corresponds to increased anxiety-like behavior after ethanol withdrawal. The unshaded region corresponds to p-values greater than 0.1, the gray-shaded region to p-values between 0.1 and 0.05, the dark gray-shaded region corresponds to p-values between 0.05 and 0.01, and the region outside of the funnel corresponds to p-values below 0.01.

### 3.2. Effects of EtOH withdrawal on anxiety-like behavior in zebrafish

Withdrawal increased time on white (Z = 2.1207, *p* = 0.0261; SMD_UB_ = -0.995; observed power = 0.6414; Figure 2A), without affecting entry duration (Z = 0.92305, *p* = 0.4301; SMD_UB_ = - 0.392; observed power = 0.147; Figure 2B). Withdrawal did not increase erratic swimming (Z = 1.9389, *p* = 0.0564, SMD_UB_ = 0.890; observed power = 0.5467; Figure 2C), but it increased risk assessment (Z = 1.9895, *p* = 0.0405, SMD_UB_ = 0.918; observed power = 0.5724; Figure 2D). Freezing (Z = 0.8082, *p* = 0.7003, SMD_UB_ = 0.286; observed power = 0.098; Figure 2E) and thigmotaxis (Z = 0.41291, *p* = 0.7024, SMD_UB_ = -0.2133; observed power = 0.0718; Figure 2F) were unaffected. As for motor effects, transitions to white were not affected by withdrawal (Z = 0.91765, *p* = 0.3763, SMD_UB_= 0.39; observed power = 0.1469; Figure 2G), but total locomotion was higher in animals exposed to withdrawal (Z = 2.5965, *p* = 0.0034, SMD_UB_= 1.31; observed power = 0.863; Figure 2H).

**Figure 2.**
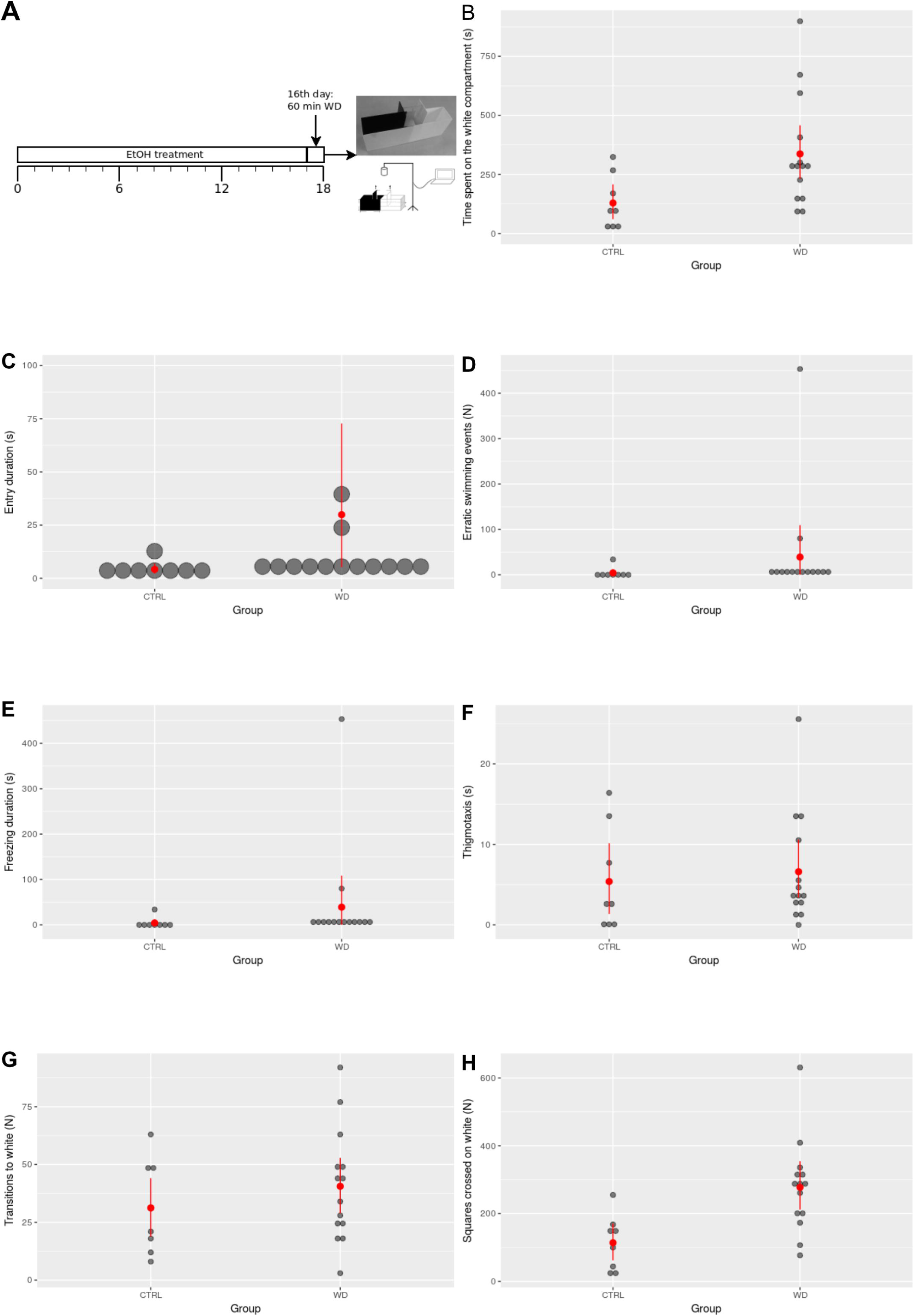
Effects of EtOH withdrawal (increasing concentration of up to 0.5% for 16 days, followed by 1 h withdrawal) on behavior in the light/dark test. (A) Scototaxis; (B) Erratic swimming; (C) Risk assessment; (D) Freezing; (E) Thigmotaxis; (F) Transitions to white; (G) Locomotion on white. Red dots represent mean, and red error bars represent nonparametric bootstrapped confidence intervals for the mean at the 95% level. To facilitate visualization, data points were jittered; therefore, their absolute position does not reflect the actual value, and values can appear to be below 0.

To control for effects of chronic exposure, independent groups were chronically exposed to EtOH (0.5%), and tested in the light/dark test immediately after the last exposure (i.e., without withdrawal). Animals exposed to EtOH spent more time in the white compartment (Z = - 1.9535. *p* = 0.0467; SMD_UB_ = -1.033; observed power = 0.482; Figure S3A), but did not show changes in entry duration (Z = -1.264, *p* = 0.2632; SMD_UB_ = 0.611; observed power = 0.204; Figure S3B). Erratic swimming was decreased by chronic EtOH treatment (Z = 2.521, *p* = 0.01; SMD_UB_ = 1.516; observed power = 0.803; Figure S3C), but no changes were observed on risk assessment (Z = 0.478, *p* = 0.793; SMD_UB_ = 0.212; observed power = 0.058; Figure S3D) or freezing (Z = 0.393, *p* = 0.736; SMD_UB_ = 0.180; observed power = 0.052; Figure S3E). Thigmotaxis was decreased by chronic EtOH (Z = 2.2879, *p* = 0.012; SMD_UB_ = 1.295; observed power = 0.671; Figure S3F). Neither transitions to white (Z = -1.790, *p* = 0.074; SMD_UB_ = -0.922; observed power = 0.402; Figure S3G) nor total locomotion (Z = -1.643, *p* = 0.097; SMD_UB_ = -0.829; observed power = 0.336; Figure S3H) were changed by chronic treatment with EtOH.

### 3.3. Effects of EtOH withdrawal on epileptic seizure-like behavior susceptibility

Only one animal from the control group exhibited Score IV epileptic seizure-like behaviors after 150 mg/kg pilocarpine, replicating findings from Pinto (2015); therefore, this dose is indeed sub-convulsive in zebrafish. All animals from the withdrawal group exhibited Score IV epileptic seizure-like behaviors after 150 mg/kg pilocarpine; after applying a log-rank model for the differences in latencies to Score IV, a significant difference was seen in the withdrawal group (χ²_[df = 1]_ = 8.8, *p* = 0.00308; Figure 3). This data can be found in our GitHub repository (https://github.com/lanec-unifesspa/etoh-withdrawal/tree/master/seizure).

**Figure 3.**
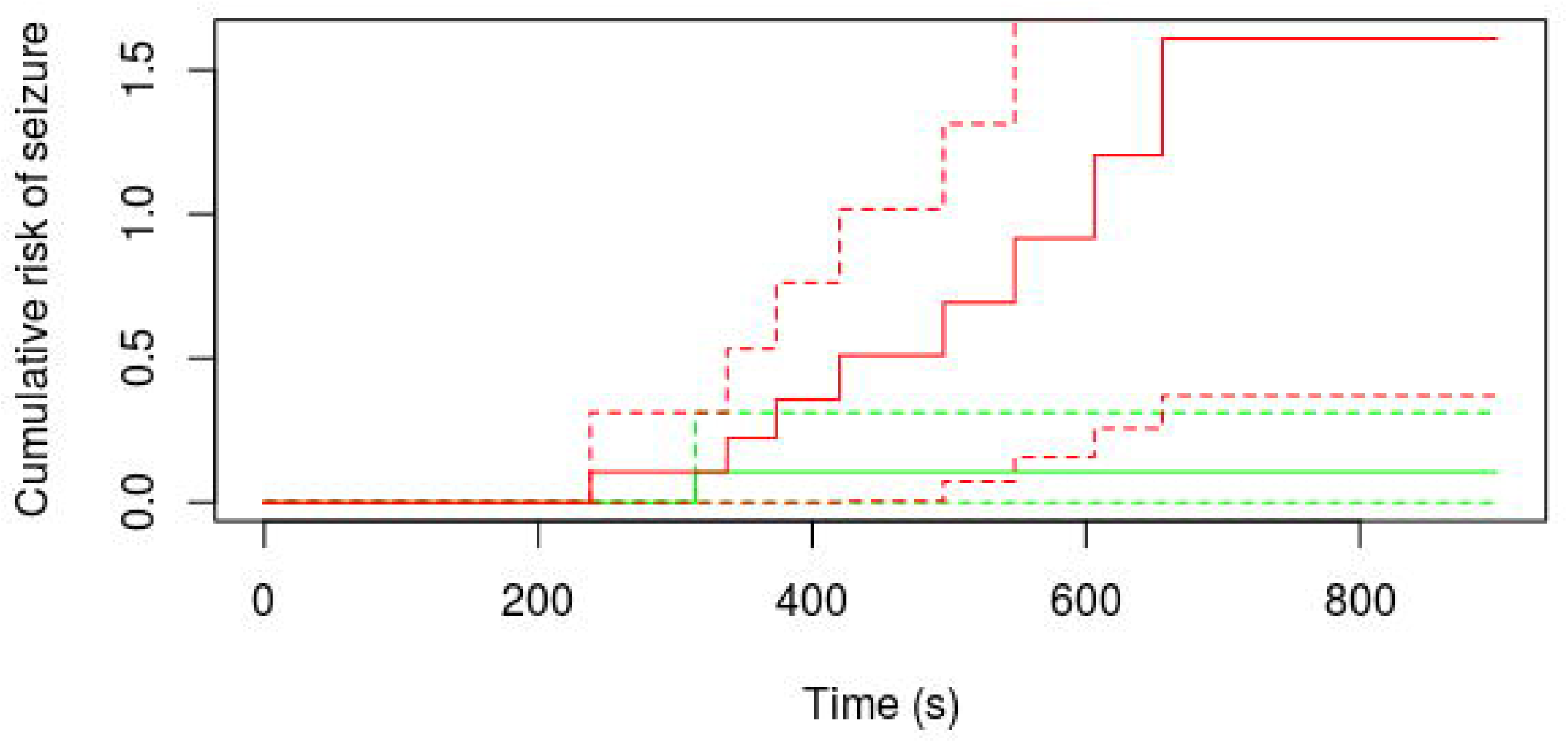
Effects of EtOH withdrawal (increasing concentration of up to 0.5% for 16 days, followed by 1 h withdrawal) on Score IV seizure latencies after sub-convulsive pilocarpine injection. Animals were observed after injection of 150 mg/kg pilocarpine in controls (green lines) and withdrawal (red lines). The dashed lines represent 95% confidence intervals around the Kaplan-Meier estimates of seizure probability at each time interval.

### 3.4. Effects of EtOH withdrawal on catalase activity

Catalase activity was reduced in the brain of animals exposed to the withdrawal regime (Z = 2.0885, *p* = 0.0156; Figure 4A), while no significant differences were found in the head kidney (Z = 1.2547, *p* = 0.1863; Figure 4B). Müller et al. (2017) also observed decreased catalase activity in the brains of zebrafish after 24 h EtOH withdrawal. Data and scripts for this experiment are available at https://github.com/lanec-unifesspa/etoh-withdrawal/tree/master/catalase.

**Figure 4.**
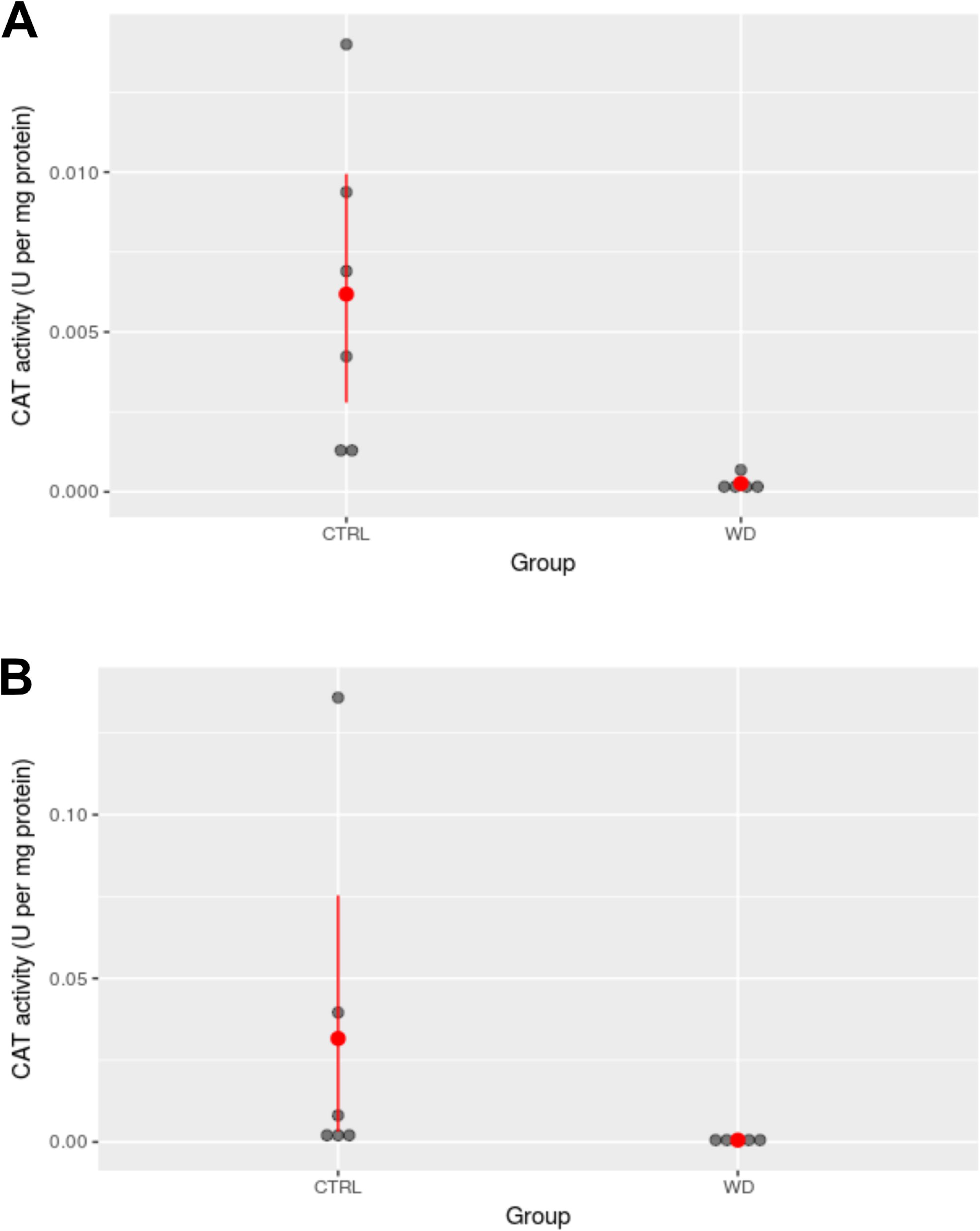
Effects of EtOH withdrawal (increasing concentration of up to 0.5% for 16 days, followed by 1 h withdrawal) on catalase activity. Enzyme activity was assessed in (A) brain and (B) head kidney of zebrafish from control (CTRL) and withdrawal (WD) groups. Red dots represent mean, and red error bars represent nonparametric bootstrapped confidence intervals for the mean at the 95% level. To facilitate visualization, data points were jittered; therefore, their absolute position does not reflect the actual value, and values can appear to be below 0.

## 4. Discussion

The present work reinforced the utility of using zebrafish as a model organism in studying ethanol withdrawal by searching for broader patterns in the literature, providing a conceptual replication of some findings regarding anxiety-like behavior and catalase activity, as well as expanding the range of behavioral domains for study. We found that the literature is inconsistent in what regards the effects of ethanol withdrawal. This heterogeneity is associated with the great procedure differences which are reported; the results of the metanalysis suggest that the main driving factors are ethanol concentration during exposure and exposure duration, with lower concentrations and longer durations more likely to induce anxiety-like behavior. Moreover, we found that ethanol withdrawal (after exposing animals for 16 days to 0.5% ethanol and 1 h withdrawal) decreased scototaxis, but increased risk assessment, in the light/dark test, and decreased the threshold for chemically-induced epileptic seizure-like behavior.

Zebrafish is increasingly being considered as a model organism in behavioral research (Bonan & Norton, 2015; Kalueff et al., 2012; Norton & Bally-Cuif, 2010; Stewart et al., 2015), with a great deal of studies on alcohol (Tran et al., 2016). Behavioral effects of drug withdrawal have been demonstrated with different drugs, including cocaine (López-Patiño, Yu, Cabral, et al., 2008; López-Patiño, Yu, Yamamoto, et al., 2008) and morphine (Cachat et al., 2010; Khor et al., 2011; Wong et al., 2010); in all cases, anxiety-like behavior was assessed. In the same direction, ethanol withdrawal has been studied mainly with models for anxiety-like behavior (Table 3). Our metanalysis revealed a significant effect of EtOH withdrawal on anxiety-like or defensive behavior in zebrafish, but a high degree of heterogeneity. Most of the heterogeneity was explained by procedural aspects; significant effects of exposure duration and EtOH concentration were found, suggesting that longer exposure and higher concentrations are critical to induce withdrawal. Other factors, including statistical inference and lack of control factors (Gerlai, 2018), could influence the heterogeneity as well. Most of the studies were underpowered, and there was an association between observed power and effect size. Risk of bias (RoB) was very high for all studies, which usually did not report blinding and random allocation; nonetheless, study quality (i.e., RoB) did not influence the results of the metanalysis, evidencing the robustness of the method. While publication bias is a relevant issue in the reproducibility and replicability of zebrafish research (Gerlai, 2018), no evidence for it was found in the systematic review.

**Table 3.**
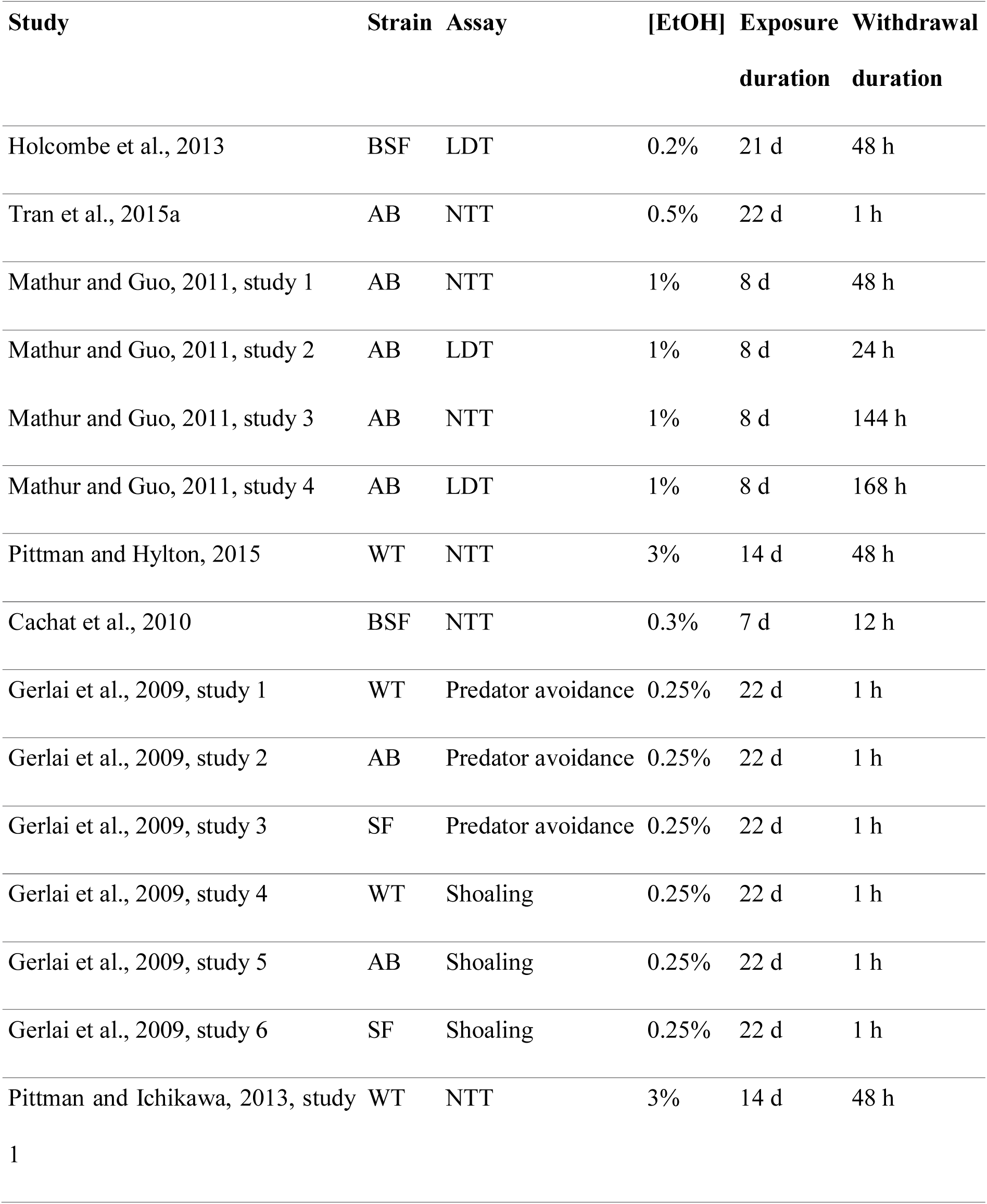
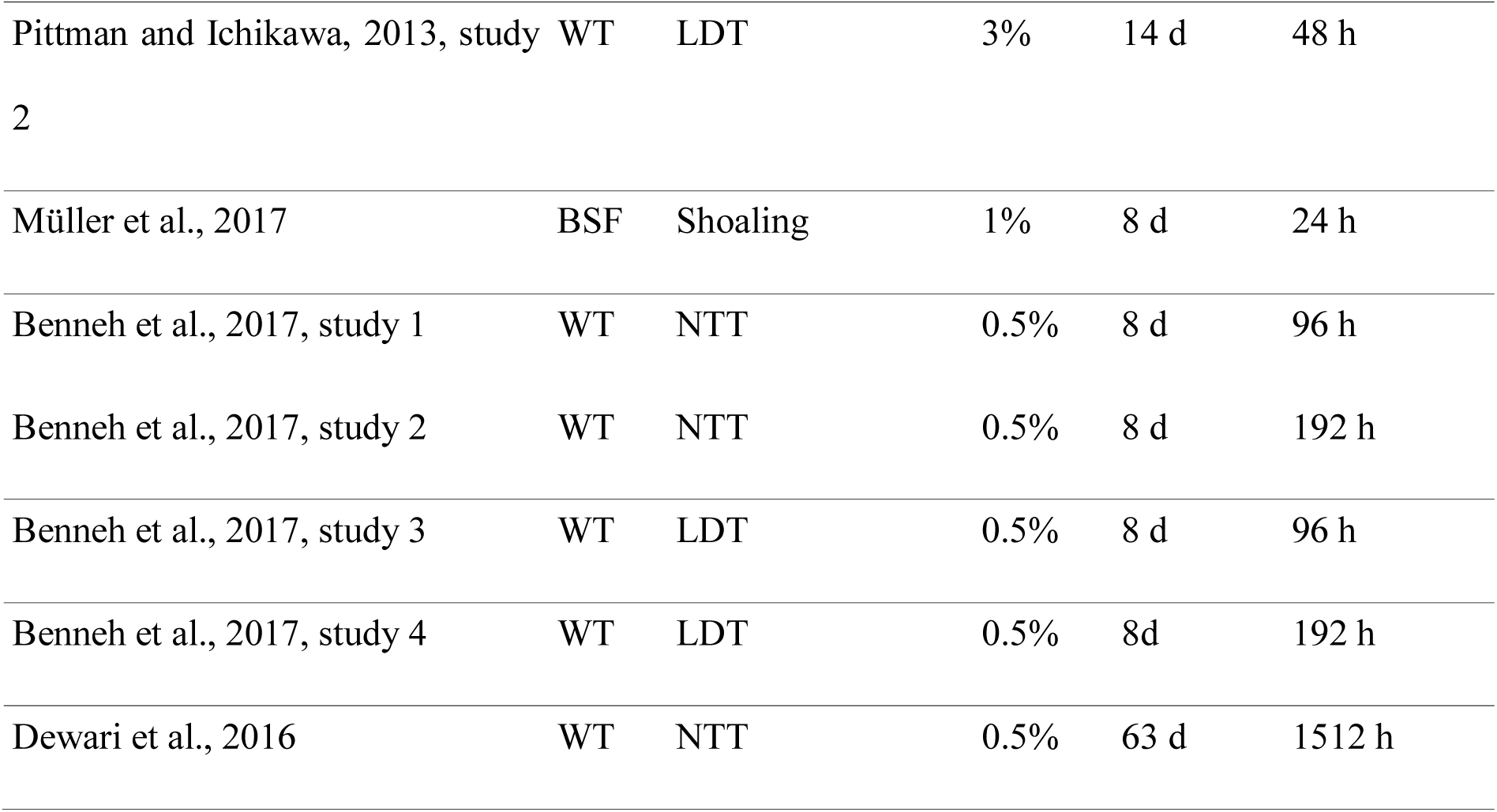
Studies included in the metanalysis. Studies were ordered by strain used (BSF = blue shortfin; WT = non-specified wild-type); behavioral assay (LDT = light/dark test; NTT = novel tank test); ethanol concentration during exposure ([EtOH]); exposure duration; and withdrawal duration.

Our conceptual replication used 60 min withdrawal, translationally relevant to the initial symptoms of EtOH withdrawal in humans (which include anxiety symptoms and epileptic seizures; Trevisan et al., 1998). Using the exposure method described in Gerlai et al. (2009), we showed that withdrawal reduces scototaxis. This is in line with the “daily-moderate” condition in Holcombe et al. (2013), which showed a decrease in scototaxis after using the same exposure profile we presented. As can be deprehended from the meta-analysis, the effect sizes calculated for scototaxis in Holcombe et al.’s (2013) experiment are similar in direction and magnitude. The Mathur and Guo (2011) scototaxis study used a higher concentration of EtOH (1%) but a shorter exposure period (8 days). While Pittman and Ichikawa (2013) used a similar exposure period (14 days), the concentration was much higher (3%); in both studies, EtOH withdrawal did not affect scototaxis.

While following only effects on scototaxis confirms findings reported in Holcombe et al. (2013), these findings contradict the overall results from the meta-analysis, which suggested that EtOH withdrawal increases anxiety-like behavior in zebrafish. If other variables are considered, however, the picture changes, with increases in risk assessment and number of transitions. The increase in risk assessment is consistent with increased anxiety-like behavior (Maximino et al., 2014), but, since scototaxis was decreased, this conclusion is only provisional. These results are difficult to interpret, but could be explained by the increased risk assessment; however, an exploratory analysis suggests negative correlation between risk assessment and time on white in the withdrawal group (r² = -0.443, vs. -0.122 in the control group). The increase in transitions without an apparent increase in locomotion in the white compartment could be interpreted as psychomotor agitation, an important symptom of EtOH withdrawal.

A reduction in scototaxis would be expected of animals chronically exposed to EtOH, which could decrease anxiety levels, and therefore not be a direct effect of withdrawal. While initially unplanned, we performed additional experiments to test this hypothesis by exposing a different group of animals to chronic ethanol and analyzing its behavior without withdrawal. We observed decreased scototaxis, but we also observed decreased erratic swimming and thigmotaxis. These results present preliminary evidence that withdrawal itself, and not the chronic exposure to EtOH, produced the reported behavioral effects.

In addition to this behavioral profile, EtOH withdrawal was also shown to decrease the threshold for chemically-induced epileptic seizure-like behavior in zebrafish. Pilocarpine is a muscarinic agonist which induces epileptic seizure-like behavior in rodents (Scorza et al., 2009), and has recently been shown to induce a similar profile in zebrafish (Pinto, 2015). The rationale of using a sub-convulsive dose is that, if EtOH withdrawal increases susceptibility to epileptic seizure-like behavior, zebrafish should present epileptic seizure-like behavior with a dose which does not induce this state in control animals. We observed increased probability of entering Stage 4 epileptic seizure-like behaviors in EtOH withdrawal animals, and a shorter latency to this event, suggesting that the protocol presented here is able to model susceptibility to epileptic seizure-like behavior, increasing the range of endpoints for studying EtOH withdrawal in zebrafish.

Finally, we also observed decreased catalase activity in the brain, but not in the head kidney of EtOH withdrawal animals. Catalase is a detoxifying enzyme that catalyzes the transformation of hydrogen peroxide, a free radical, into water and oxygen (Aebi, 1984); in the brain, catalase is poorly expressed, but usually associated with microglial activity (Dringen, 2005). The inhibition of enzymatic activity observed here replicates results by Müller et al. (2017) in the zebrafish brain; in that study, however, the exposure duration was shorter (8 days), the concentration of EtOH was higher (1%), and the pattern of exposure was different (animals were exposed for 20 min per day, instead of continuously, to ethanol) compared to the present investigation. The lack of effect on the head kidney, where interrenal cells (the teleost functional equivalent of the mammalian adrenal cortex) lie, suggesting that possible effects on cortisol (e.g., Cachat et al., 2010) are not due to effects in these cells, but upstream in the hypothalamus-hypophyseal-interrenal axis.

The present work contributed to the use of zebrafish as a model in EtOH withdrawal research by: A) identifying sources of heterogeneity in the literature on EtOH withdrawal and anxiety-like behavior in the species; B) presenting a conceptual replication of withdrawal-induced anxiogenesis in the light/dark test; C) extending the behavioral phenotypes to include epileptic seizure-like behavior susceptibility; and D) replicating the effects on catalase activity, suggesting that EtOH withdrawal-elicited oxidative stress could be a mechanism of anxiogenesis in the species. Further work will characterize these mechanisms with care.

## Acknowledgements

SNSC was the recipient of a CNPq/PIBIC studentship. GRF was the recipient of a CNPq/PIBITI studentship. WEAO was the recipient of a CNPq/PIBIC-EM studentship to high school students. This manuscript appeared as a preprint on bioRxiv (doi: 10.1101/201368).

---

**Figure S1** Influential study analysis for the metanalysis. Statistics represent standardized residuals (rstudent), DFFITS (dffits), Cook’s distances (cook.d), covariance ratios (cov.r), estimates of τ*²* (tau2.del) and test statistics (QE.del) for (residual) heterogeneity when each study is removed in turn, hat values, and weights for each of the 16 studies examining the effects of EtOH withdrawal on zebrafish defensive behavior. Red filling indicates an influential study.

**Figure S2** Relationship between observed power and modulus SMD values in the metanalysis. Curve fitting was made using a linear model.

**Figure S3** Effects of chronic ethanol exposure on behavior in the light/dark test. (A) Scototaxis; (B) Erratic swimming; (C) Risk assessment; (D) Freezing; (E) Thigmotaxis; (F) Transitions to white; (G) Locomotion on white. Red dots represent mean, and red error bars represent nonparametric bootstrapped confidence intervals for the mean at the 95% level. To facilitate visualization, data points were jittered; therefore, their absolute position does not reflect the actual value, and values can appear to be below 0.

