## Supplementary figures and images for "Behavioral and biochemical effects of ethanol withdrawal in zebrafish"

### Supplementary Materials

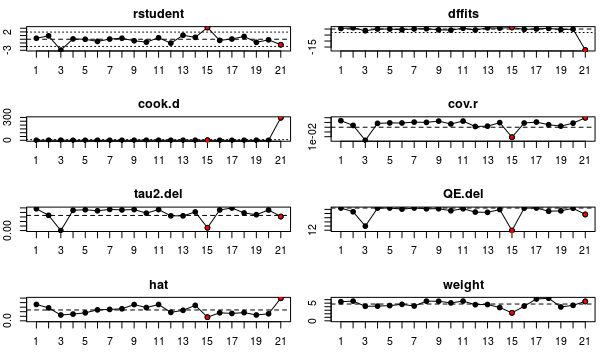

### Supplementary Materials

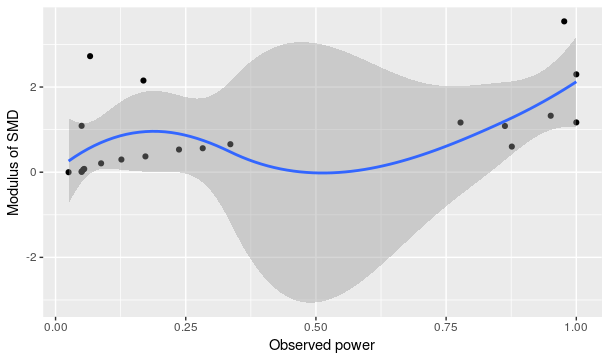

### Supplementary Materials

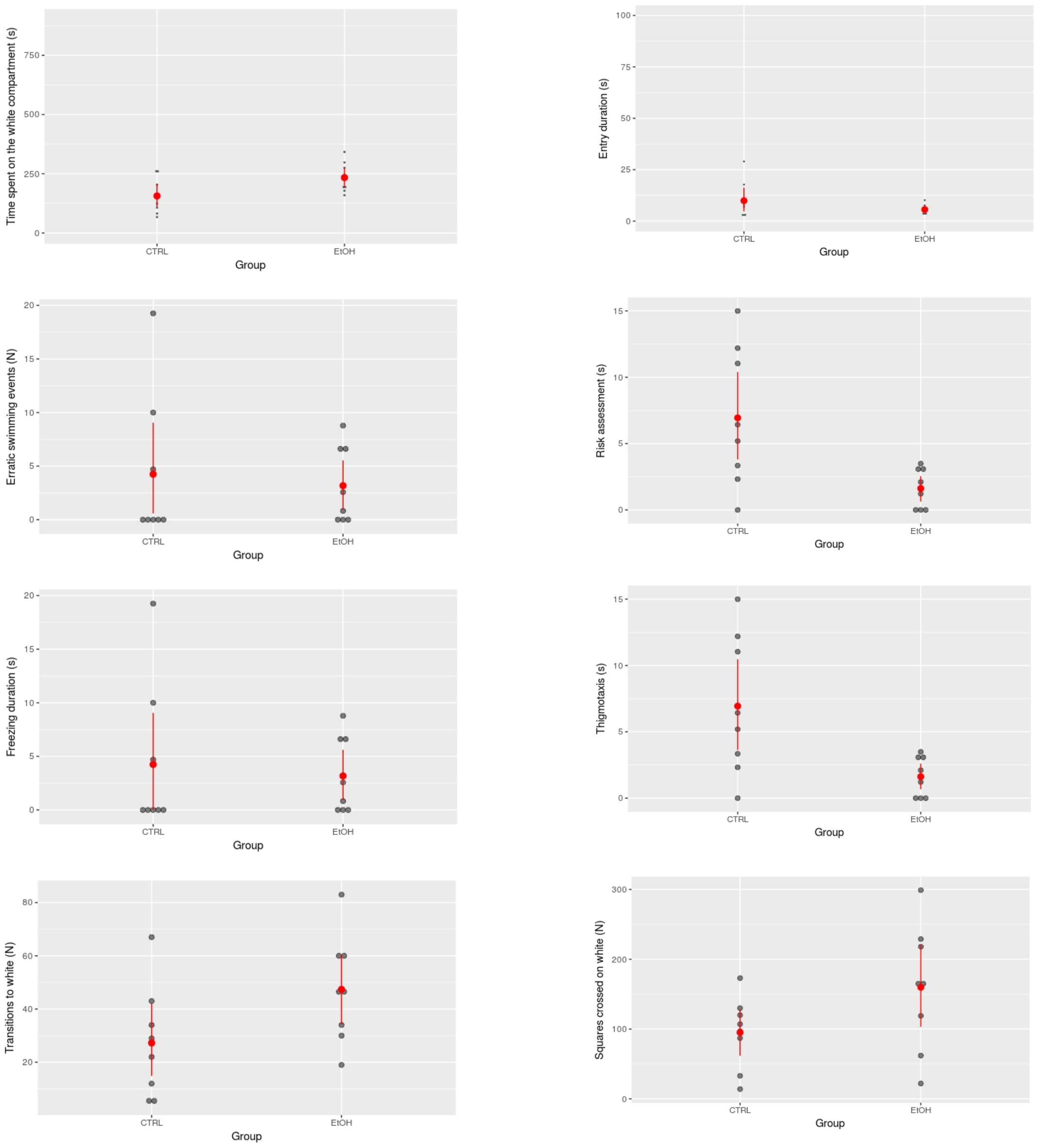
